# Intestinal microbiota modulates the anti-tumor effect of oncolytic virus vaccine in colorectal cancer

**DOI:** 10.1101/2023.05.28.542655

**Authors:** Xia Chen, Guang-Jun Wang, Ling Qin, Bing Hu, Jun Li

**Affiliations:** Department of Gastroenterology and Hepatology, West China Hospital, Sichuan University, Chengdu, 610041, China; Department of Gastroenterology, Clinical medical college and the first affiliated hospital of Chengdu medical college, Chengdu, 610500, China

**Keywords:** Colorectal cancer, Immunotherapy, Intestinal microbiota, Oncolytic virus vaccine.

## Abstract

**Background:** Immunotherapy such as oncolytic virus has become a powerful cancer treatment but only a part of cancer patients can benefit from it, especially to advanced-stage cancer patients are required new therapeutic strategies to facilitate extended survival. Intestinal microbiota may contribute to colorectal cancer (CRC) carcinogenesis and response to immunotherapy. However, whether and how the modulating effect of intestinal microbiota on oncolytic virus vaccine (OVV) in CRC remains to be investigated.

**Methods:** We generated a MC38-gp33 CRC mouse model and treated with OVV-gp33 in early- and advanced-stages. Probiotics, fecal microbiota transplantation (FMT) and antibiotics (ABX) were treated to regulate the microbial composition of CRC mice of advanced stage. The tumor growth rate and survival time of mice were recorded. 16S rDNA sequencing analyzed the microbial composition and flow cytometry detected the T cells subsets activity.

**Results:** OVV-gp33 treatment led to inhibited tumor growth and prolonged survival in the early stage of CRC but did not have a significant effect on the advanced stage of CRC. Moreover, 16S rDNA sequence analysis and flow cytometry showed significant differences in intestinal microbiota composition, microbial metabolites and T-cell subsets in early- and advanced-stage CRC. Probiotic and FMT treatment significantly enhanced the antitumor effect of OVV in advanced stage of CRC with an increased abundance of activated CD8^+^ T cells and a decreased ratio of Treg cells, while depletion of the microbiota by ABX eliminated the antitumor activity of OVV with decreased CD8^+^ T-cell activation and upregulated Treg cells.

**Conclusions:** These results indicate that intestinal microbiota and microbial metabolites play an important role in the OVV antitumor effect in CRC, furthermore, altering the intestinal microbiota composition can modulate the antitumor and immunomodulatory effect of OVV in CRC.

## Introduction

Colorectal cancer(CRC)is one of the most common malignant tumor in the world, contributes to the second leading cause of cancer related deaths[1]. It is estimated that the incidence of CRC would increase to more than 2.2 million new cases and 1.1 million deaths by 2030 [2]. The global incidence varies significantly between different regions therefore screening programs are still need to be implemented especially in moderate- to high-incidence countries[3]. The survival rate of CRC is closely related to different clinical stages, which ranges from greater than 90% in CRC patients with stage I to slightly greater than 10% in patients with stage IV[4]. Therefore, it is necessary to explore new therapeutic strategies to facilitate extended survival especially in the middle - and advanced - stage CRC patients.

Immunotherapy to control or modulate the immune system has emerged as a powerful clinical strategy for cancer treatment[5]. Additionally, immunotherapy alone or combination with other existing therapies providing opportunities to improve clinical outcome of advanced cancer. However, these therapeutics apply to clinical use faces several challenges related to both efficacy and safety[6]. Currently, considerable interest raised in other novel strategies that might have promoted antitumor benefit through the use of complementary agents without overlapping toxicity characteristics, such as oncolytic virus (OV)[7].OV is a virus with the ability to replicate, selectively expand and kill tumor cells without affecting healthy cells. In addition to killing infected tumor cells, OV can also augment anti-tumor immune response by lysing tumor cells to release pathogen-related molecules and tumor-related antigens, activate and recruit specific T cells, thus display antitumor effect against cancer[8, 9]. Oncolytic virus vaccine (OVV) is a remolding of OV, which further enhances the immune response and clinical efficacy against tumors by targeting tumor-specific antigens[7]. The application of OVV has been reported to successfully enhance cancer suppression and eradication in solid tumors[10, 11]. However, far too little attention has been paid to OVV in the treatment of CRC. Given its potential role in shaping immunity, we can anticipate that targeted antigen of OVV directed to CRC will become an integral part of cancer treatment.

The interactions between intestinal microbiota and tumors have attracted much attention in modulating the efficacy and toxicity of anticancer drugs including traditional chemotherapy and the recently successful immunotherapy [12, 13]. Intestinal microbiota, as an important component of the tumor microenvironment, influences disease in regulating not only mucosal but also systemic immune responses[14]. Previous studies have showed numerous microbiotas being altered abundance in CRC patients, which may enhance CRC antitumor immunity and directly contribute to development of chemoresistance, carcinogenesis and tumor progression [15, 16]. Approaches that modulate the composition of intestinal microbiota may offer a promising strategy for the treatment of CRC. However, whether and how intestinal microbiota affect the therapeutic effect of OVV in CRC has not been investigated before. Here, we investigated the antitumor effect of OVV and the microbiota composition in different stages of CRC. On the other hand, we characterized how microbiota composition alteration influenced the therapeutic effect of OVV in CRC and the subsequent T-cells activity.

## Methods and materials

### Mice

C57BL/6 mice were all female, 4-5 weeks in age and purchased from Chengdu Dashuo Animal Co., Ltd. Mice were housed under standard conditions at 24±2 °C with food and water provided.

### Ethics statement

All experimental protocols were approved by the animal experimental teaching and research committee of Chengdu medical college (approval no. 2017009), and were performed in accordance with the nation’s relevant laws and animal welfare. All efforts were made to minimize animal suffering.

### Cell lines and Culture Conditions

MC38 colon cancer cells were purchased from ATCC. These cell lines were authenticated by STR profiling. Cells within three to eight passages were cultured in complete Dulbecco’s Modified Eagle Medium (DMEM; Thermo Fisher Scientific) containing 10% FBS (Thermo Fisher Scientific) and maintained at 37 °C with 5% CO_2_. MC38 cells were seeded in a 24-well plate (100,000 cells/well), and MC38-gp33 cells were generated as described previously[17].

### CRC Mouse Model and grouping

Mice were divided into 6 groups (5 mice per group). Mice that did not receive tumor implantation were used in the control group (CON). CRC mice were implanted subcutaneously (s.c.) with 10^7^ MC38-gp33 cells in 20 µl PBS. Mice implanted with MC38-gp33 cells were injected with OVV-gp33 by tail vein on Day 5 (defined as the early CRC [ECRC] group) and Day 17 (defined as the advanced CRC [ACRC] group) post-tumor inoculation. To deplete the microbiota in the antibiotics (ABX) group, mice received ABX (1 g/L ampicillin, 160 mg/L neomycin sulfate, 1 g/L metronidazole, 0.5 g/L vancomycin) solution for 2 weeks before tumor inoculation, followed by continued ABX water application for 17 days and administration of OVV-gp33 on Day 17 after tumor inoculation. CRC mice in the probiotic (PRO) group were gavaged with solutions including *Bifidobacterium longum*, *Lactobacillus bulgaricus* and *Streptococcus thermophilus* (100 mg/ml, 0.7 g/kg/d) once every 24 h for 17 consecutive days and then injected with OVV-gp33 on Day 17. For the fecal microbiota transplantation (FMT) group, FMT was performed on the donor stool of age-matched healthy C57BL/6 female mice every other day after tumor inoculation until OVV-gp33 administration on Day 17.

Tumor volume was measured every 5 days using calipers and calculated as (x × y × z)/ mm^3^. Mice with the tumor reaching 1000mm^3^ or with > 10% body weight or ulcerated tumor were euthanized before the end of the study. Mice were euthanized by CO_2_ inhalation followed by cervical dislocation.

### Fecal microbiota analysis

The fecal samples of mice in ECRC were collected at Day 5 post-tumor inoculation before OVV-gp33 treatment, while the fecal samples of other CRC mice were collected at Day 17 post-tumor inoculation but before OVV-gp33 treatment. The fecal samples for 16S rDNA sequencing were frozen at -80 °C and then delivered to Genesky Biotechnologies Inc. (www.geneskybiotech.com/) on dry ice for further 16S rDNA sequencing. Next, alpha diversity, beta diversity and intestinal microbial species composition were analyzed for every sample. Linear discriminant analysis effect size (LEfSe) was used to select significantly enriched candidates at the phylum and genus levels.

### Flow cytometry

Mice were euthanized, and blood samples were taken from the orbital venous plexus. Briefly, 100 μl blood samples were added to 1 ml RBC lysis buffer (Biosharp) for 2 min at 4 °C and then centrifuged at 300 × g for 5 min. The cell precipitates were washed with 3 ml PBS 2 times and suspended in 200 μl flow buffer. The cell suspension was first stained with 1 µl of Fixable Viability Stain 780 for 15 min. After washing with flow buffer, single cells were incubated with 5 μl CD8^+^ antibody (BD Pharmingen), 2 μl of CD3 antibody (BD Pharmingen), 5 μl of CD4 antibody (BD Pharmingen) and 43 μl of flow buffer for 30 min at 4 ℃. After washing the cells with flow buffer, the cells were fixed and permeabilized using the FoxP3 staining Buffer Set (BD Pharmingen) at 4 ℃ according to the manufacturer’s protocol. Then, the cells were incubated with 5 μl of FoxP3 antibody (BD Pharmingen) for 40 min, and flow cytometry data were acquired.

### Statistical Analysis

The statistical analysis of the data was performed using SPSS, version 25.0 and GraphPad Prism, version 9.0. The data are expressed as the means ± SD. The results of different treatment groups were compared by one-way analysis of variance. The numerical variables between 2 groups were compared using Student’s *t* test. Differences were considered significant with *P* values < 0.05.

## Results

### OVV achieved antitumor response in CRC mice by regulating T cell activity

First, we evaluated the antitumor effects of OVV-gp33-targeting therapy on MC38-gp33 bearing mice during tumor progression. We found that more than 60% of mice in ECRC group were cured. The tumor growth was inhibited significantly on the 12^nd^ day after inoculation, this effect continued to the 17^th^ day, and only less than 20% of mice were not cured completely (Fig 1A), indicating that OVV-gp33 targeted therapy had a significant therapeutic effect on early stage of CRC. In ACRC group, the tumor mass continued to grow after inoculation with tumor cells, treatment of OVV-gp33 on the 17^th^ day could only slow the growth of tumor but could not achieve the curing effect (Fig 1B). We measured the tumor volume of CRC mice on the 21^st^ day post-tumor inoculation, found that the tumor volume in ACRC group was significantly larger than that in ECRC group, and the growth rate was also faster than that in ECRC group (Fig 1C,1D). The total observation period in this study was 60 days. At approximately 40 days of observation, mice in ACRC group reached the end event, while the cured mice in ECRC group were observed for 60 days without recurrence, and the survival time of mice in ACRC group was significantly lower than that in ECRC group (Fig 1E). These results suggested that OVV application appeared to be effective in treating the early stage of CRC, and this significant curative effect started on day 12, but it was not as effective in the advanced stage of CRC.

**Fig 1.**
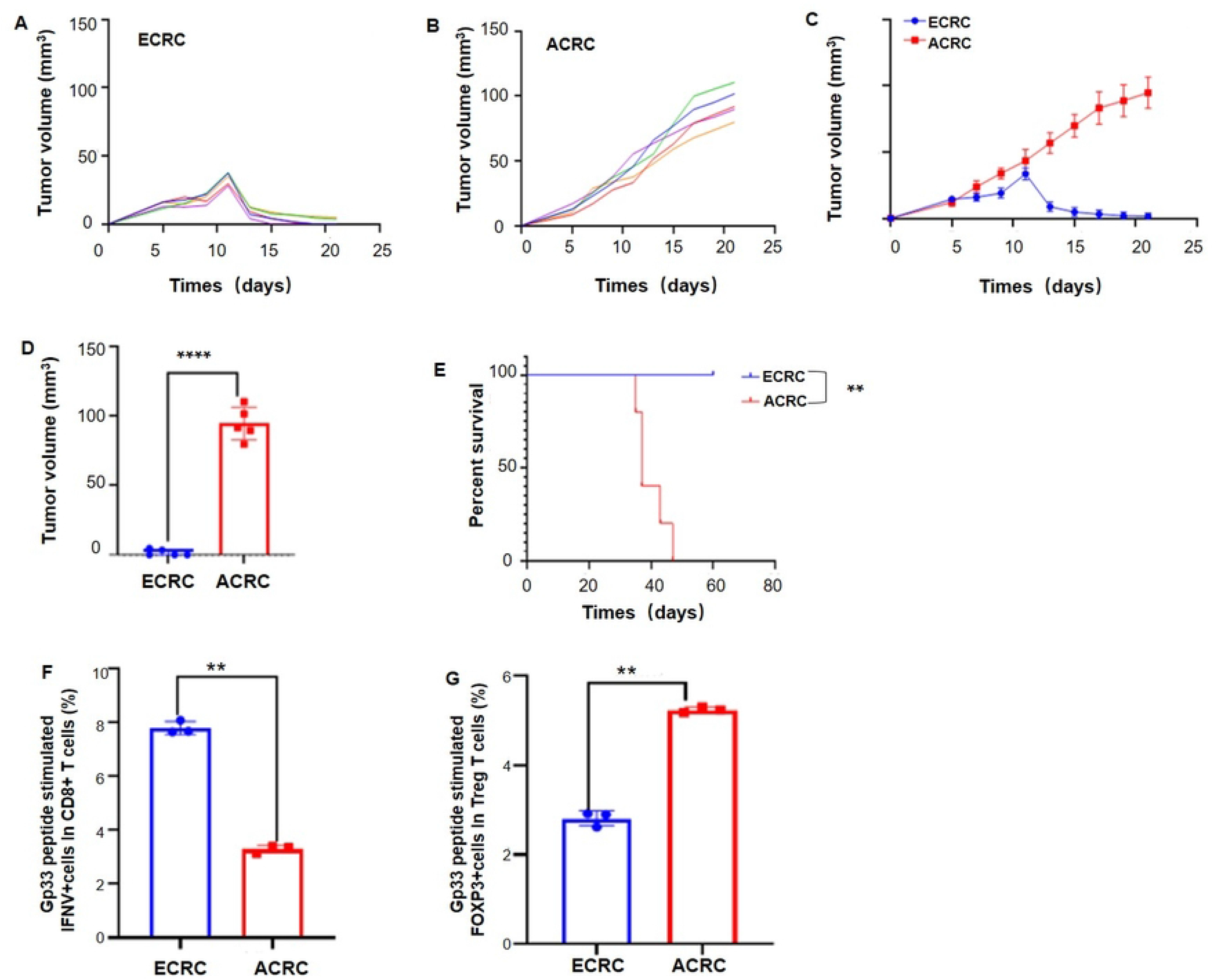
OVV achieved anti-tumor response in CRC mice by regulating T cell activity. Tumor growth curve of ECRC group (A) and ACRC group (B). (C), (D) Comparation of tumor volume between ECRC group and ACRC group. **** *P*<0.01. (E) Survival curve of ECRC group and ACRC group. ** *P* < 0.05. (F) Activity of CD8^+^ T cells. \*\**P*<0.05. (G) Ratio of Treg cells. \*\**P*<0.05.

T cells have an important role in the treatment of tumor, especially CD8^+^ T cells mediate antitumor immune response through recognition of tumor antigens and direct killing of transformed cells[18, 19]. As a heterogeneous subset of immunosuppressive T cells, regulatory T (Treg) cells characterized by expression of FOXP3, which often represents a critical barrier to anti-tumor immunity and immunotherapy[20, 21]. Flow cytometric results indicated that the abundance and activity of CD8^+^ T cells increased significantly in early-stage CRC mice compared to advanced-stage CRC mice.

Activated CD8^+^ T cells were found in 8.07% and 3.39% of mice in ECRC group and mice in ACRC group, respectively (Fig 1F). The ratio of activated Treg cells in early-stage CRC mice also differed from that in advanced-stage CRC mice, which were 2.64% and 5.31%, respectively (Fig 1G). Collectively, these results suggest that the anti-tumor effect of OVV is at least partly dependent on modified subsets of T cells.

### Intestinal microbiota influenced the therapeutic effect of OVV in CRC mice

Having shown that the time of CRC tumor formation affected the therapeutic effect of OVV-gp33, the underlying mechanism was then investigated. Previous studies showed that the intestinal microbiota plays important roles in the development of CRC and microbiota abundance is altered in CRC patients[22, 23]. We next investigated the characteristics of the intestinal flora based on fecal sample 16S rDNA sequencing. Alpha diversity analysis showed that after intradermal tumor formation, the total abundance and diversity of microbiota in CRC mice increased compared with that in mice without tumor inoculation. The abundance and diversity of the intestinal microbiota in mice of ACRC group decreased with the development of tumors, but it was still higher than that in mice without tumor inoculation (Fig 2A). Similarly, beta diversity analysis showed that the intestinal microbiota composition in CRC mice changed significantly with the progression of CRC (Fig 2B). The analysis of microbial species composition showed that, at the phylum level, Bacteroidetes, Firmicutes, Proteobacteria and Actinomyces were dominant phyla in mice of ECRC group, but with the continuous development of CRC, Bacteroidetes and Proteobacteria increased while Firmicutes decreased (Fig 2C). The main bacteria enriched in mice of ECRC group were Enterobacterium, Rumen coccus, *Psychrobacter*, *Clostridium*, *Aeromonas*, etc, while the enriched bacteria in mice of ACRC group were *Allobaculum*, *Staphylococcus* and *Marvinbryantia* (Fig 2D).

**Fig 2.**
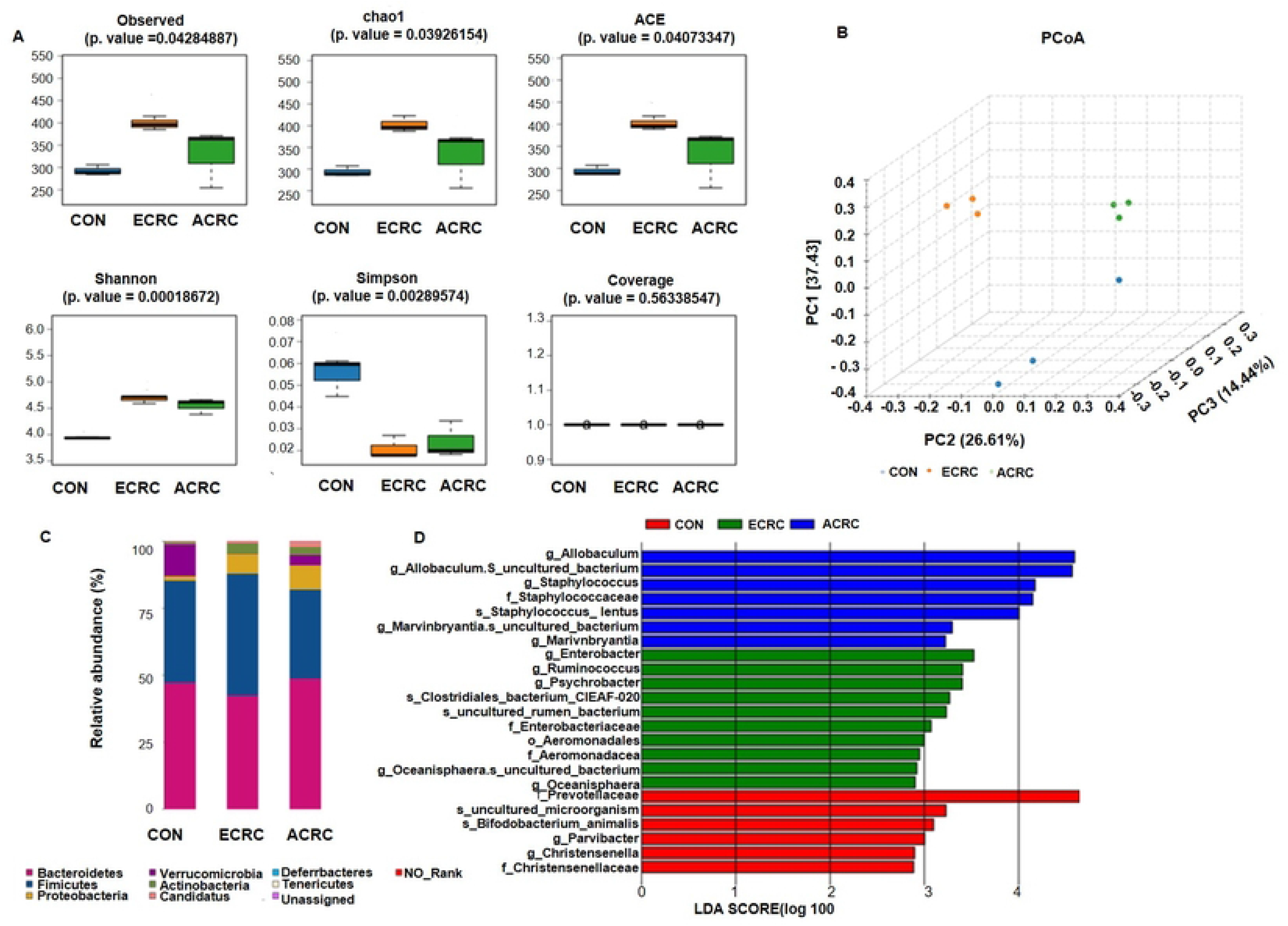
Intestinal microbial composition in early and advanced stage of CRC. (A) Alpha diversity analysis of stool samples. (B) PCoA analysis of stool samples. (C) Analysis of microbial species composition. (D) LEfSe analysis.

KEGG metabolic pathway analysis showed that, with the progression of CRC, significantly down-regulated pathways were enriched in ansamycin biosynthesis, fatty acid biosynthesis, pentose phosphate pathway, etc. Significantly up-regulated pathways are concentrated in secondary bile acid biosynthesis, D alanine metabolism, vancomycin antibiotic biosynthesis, etc (Supplementary Fig 1A). COG differential protein function analysis showed that significantly down-regulated protein functions were mainly concentrated in DNA-binding transcription regulators, hydroxymethylpyrimidine pyrophosphatases and other HAD family phosphatases, etc. Significantly upregulated differential protein functional enrichment in glycosyltransferase involved in cell wall disynthesis, multidrug efflux pump subunit AcrA (membrane fusion protein), etc (Supplementary Fig 1B).

We next used ABX to deplete the gut microbiota to further define the role of the intestinal microbiota in OVV-gp33 therapeutic efficacy. We observed an accelerated tumor growth rate in ABX-treated mice and no obvious tumor inhibition effect after OVV-gp33 treatment, which was significantly different from other advanced stages of CRC in mice without ABX depletion (Fig 3A-D). Survival analysis showed that ABX application significantly reduced the survival rate, reaching the end event on Day 25 after inoculation (Fig 3E). In addition, depletion of the intestinal microbiota by ABX prevented CD8^+^ T-cell activation and increased the ratio of Treg cell (Fig. 3F, 3G), 16S rDNA sequencing revealed that ABX reduced the abundance and diversity of the intestinal microbiota and microbial species composition, and the main enriched intestinal flora were Enterobacterium, *Escherichia*, *Morganella*, *Citrobacter*, *Morganella*, and *Thiomonas* (Fig 4A-4D). The enrichment pathway in mice of ABX group was involved in lipoic acid metabolism, folic acid biosynthesis, other glycan degradation, amino sugars and nucleotide sugars metabolism (Supplementary Fig 2A). The significantly down-regulated protein functions were mainly concentrated in aryl sulfonated enzyme A or related enzymes, multi-drug effector pump subunit AcrB, protein n-acetyltransferase, outer membrane protein, etc. Significantly upregulated differential protein function is enriched in ABC lipoprotein output system, ATPase component, DNA-binding transcription regulatory factor, etc (Supplementary Fig 2B).

**Fig 3.**
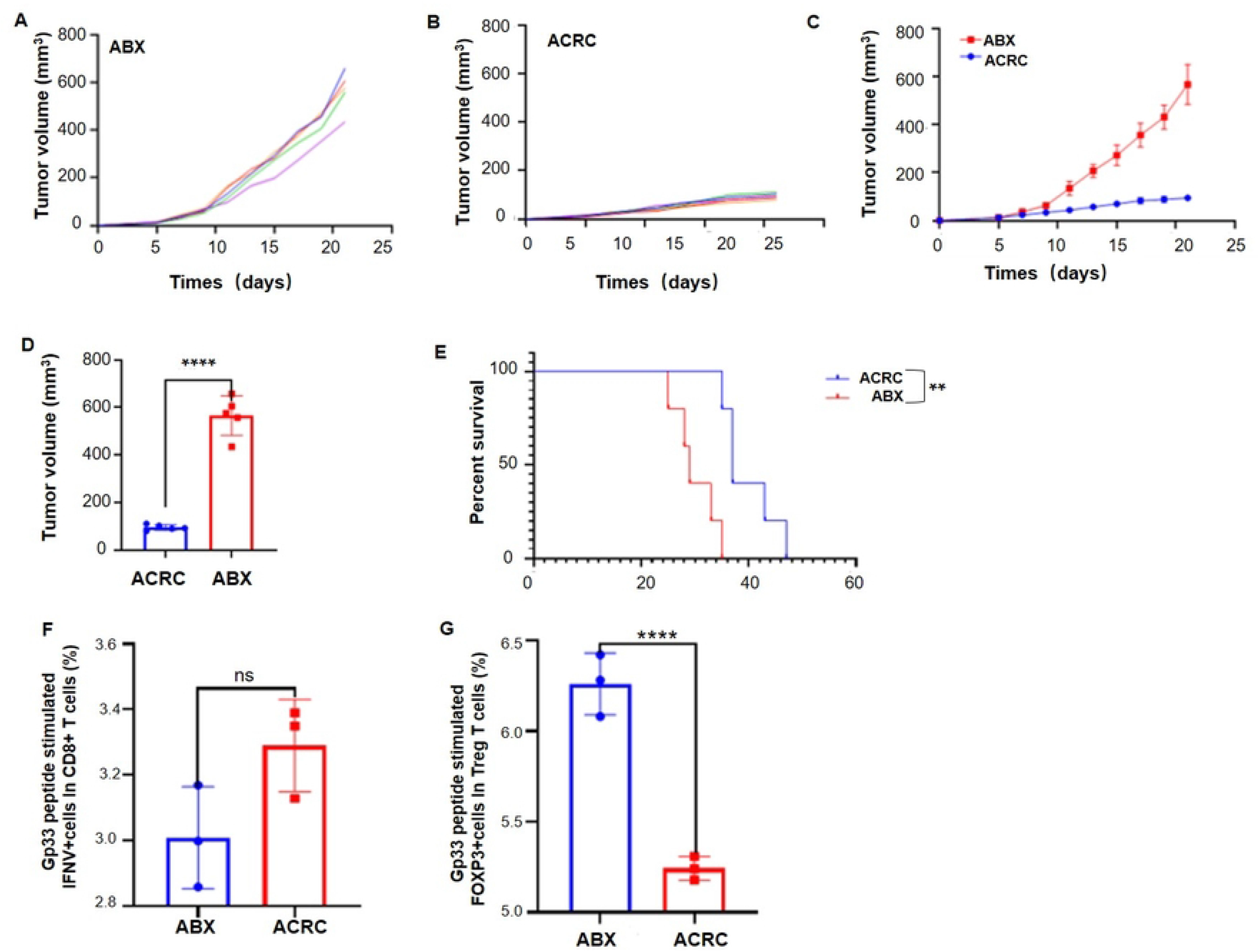
ABX accelerated tumor growth and influenced T cells activity in CRC mice. Tumor growth curve of ABX group (A) and ACRC group (B). (C), (D) Comparation of tumor volume between ABX group and ACRC group. \*\*\*\**P*<0.01. (E) Survival curve of ABX group and ACRC group. \*\**P*<0.05. (F) Activity of CD8^+^ T cells. ns: no significance. (G) Ratio of Treg cells. \*\*\*\**P*<0.01.

**Fig 4.**
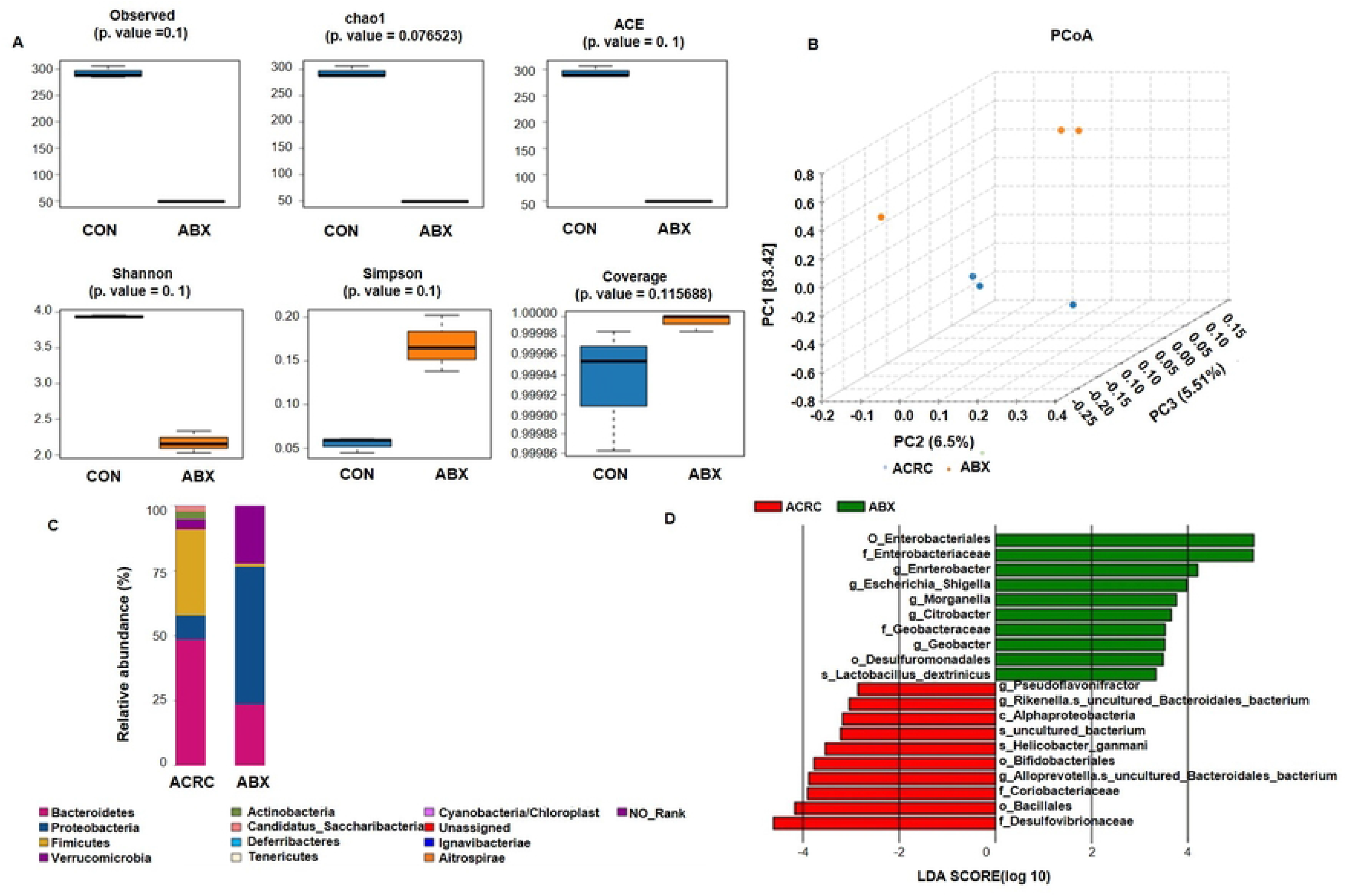
Intestinal microbial composition in ABX-treated CRC mice. (A) Alpha diversity analysis of stool samples. (B) PCoA analysis of stool samples. (C) Analysis of microbial species composition. (D) LEfSe analysis.

### Probiotic enhanced the therapeutic efficacy of OVV in advanced stage of CRC mice

In order to address the modulating effect of microbiota on the efficacy of OVV in CRC, we treated the advanced stage of CRC mice with probiotics. Mice were gavaged with probiotics for 17 days and then OVV-gp33 treated on the 17th day after inoculation. Results showed that the tumor growth rate of mice received probiotics was slower than that did not, and this tumor inhibition effect was enhanced by OVV-gp33 treatment on day 17, while the growth rate of advanced stage of CRC mice only received OVV-gp33 unchanged (Fig 5A, 5B). When CRC mice in PRO group received OVV-gp33 treated at days 17, the tumor growth slowed again, and the tumor inhibition effect enhanced than those of mice without probiotics treatment (Fig 5C). On day 21 post-tumor inoculation, the tumor mass volume of mice in ACRC group was significantly larger than that in PRO group (*P* < 0.05) (Fig 5D). Mice in ACRC group reached the end event at approximately 40 days of observation. In contrast, mice in PRO group reached the end event at approximately 50 days, and the survival time of mice in this group was longer than that in ACRC group (Fig 5E).

**Fig 5.**
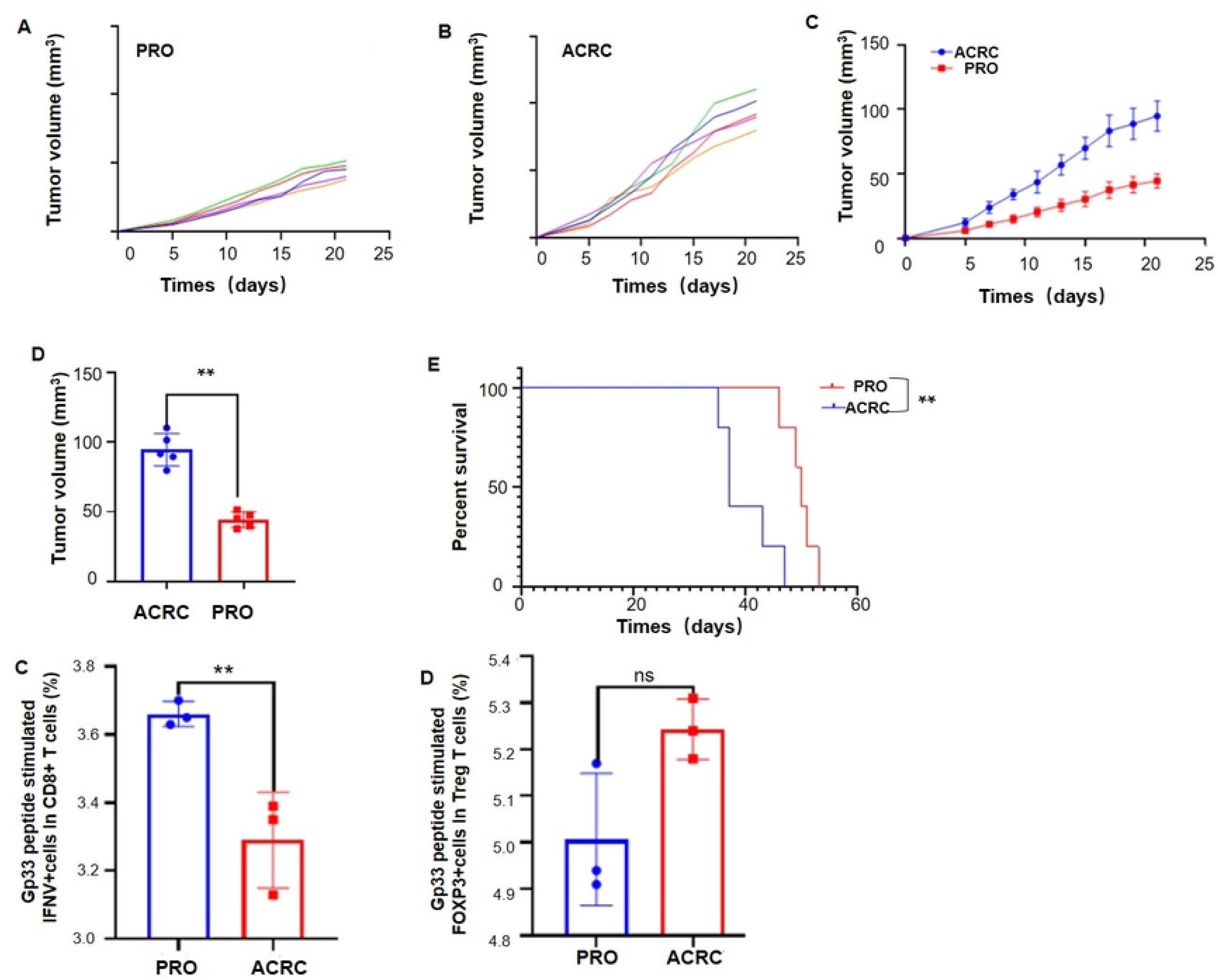
Probiotics improved the therapeutic effect of OVV-gp33 treatment. Tumor growth curve of PRO group (A) and ACRC group (B). (C), (D) Comparation of tumor volume between PRO group and ACRC group. ** *P* < 0.05. (E) Survival curve of PRO group and ACRC group. ** *P*<0.05. (F) Activity of CD8^+^ T cells. \*\**P*<0.05. (G) Ratio of Treg cells. ns: no significance.

This underscores the enhancement effect of probiotics on antitumor responses and the survival-improving effect of OVV-gp33 in advanced stage of CRC. The CRC mice treated with probiotics and OVV-gp33 showed an increase in the abundance of activated CD8^+^ T cells up to 3.63%, and the difference was significant when compared to CRC mice without probiotic treatment (*P*<0.05) (Fig 5F). In contrast, probiotic intervention decreased the ratio of Treg cells in OVV-gp33-treated CRC mice to 4.94% (Fig 5G).

16S rDNA sequencing results showed that the abundance and diversity of microbiota in mice of PRO group were lower than those in mice of ACRC group, but the difference was not statistically significant (Fig 6A). Meanwhile, beta diversity PCoA showed that the intestinal flora composition of CRC mice treated with probiotics was significantly changed (Fig 6B). The analysis of microbial species composition showed that at the phylum level, Bacteroidetes, Firmicutes, Proteobacteria and *Vermicella* were dominant in mice treated with probiotics, the amounts of Bacteroidetes and Proteobacteria were significantly reduced, and Firmicutes and Verrucomicrobia increased significantly compared with mice in ACRC group (Fig 6C). The bacteria enriched in mice treated with probiotics were *Desulfovibrio*, *Oisenella*, *Marvinbryantia*, *Escherichia*, etc (Fig 6D). The enrichment pathways of mice in PRO group were D alanine metabolism, glycolysis/gluconeogenesis, secondary bile acid biosynthesis, fatty acid biosynthesis, etc (Supplementary Figure 3A). The significantly down-regulated protein functions were mainly concentrated in amino acid transporters, DNA-binding transcription regulators, LacI/PurR family, etc.

**Fig 6.**
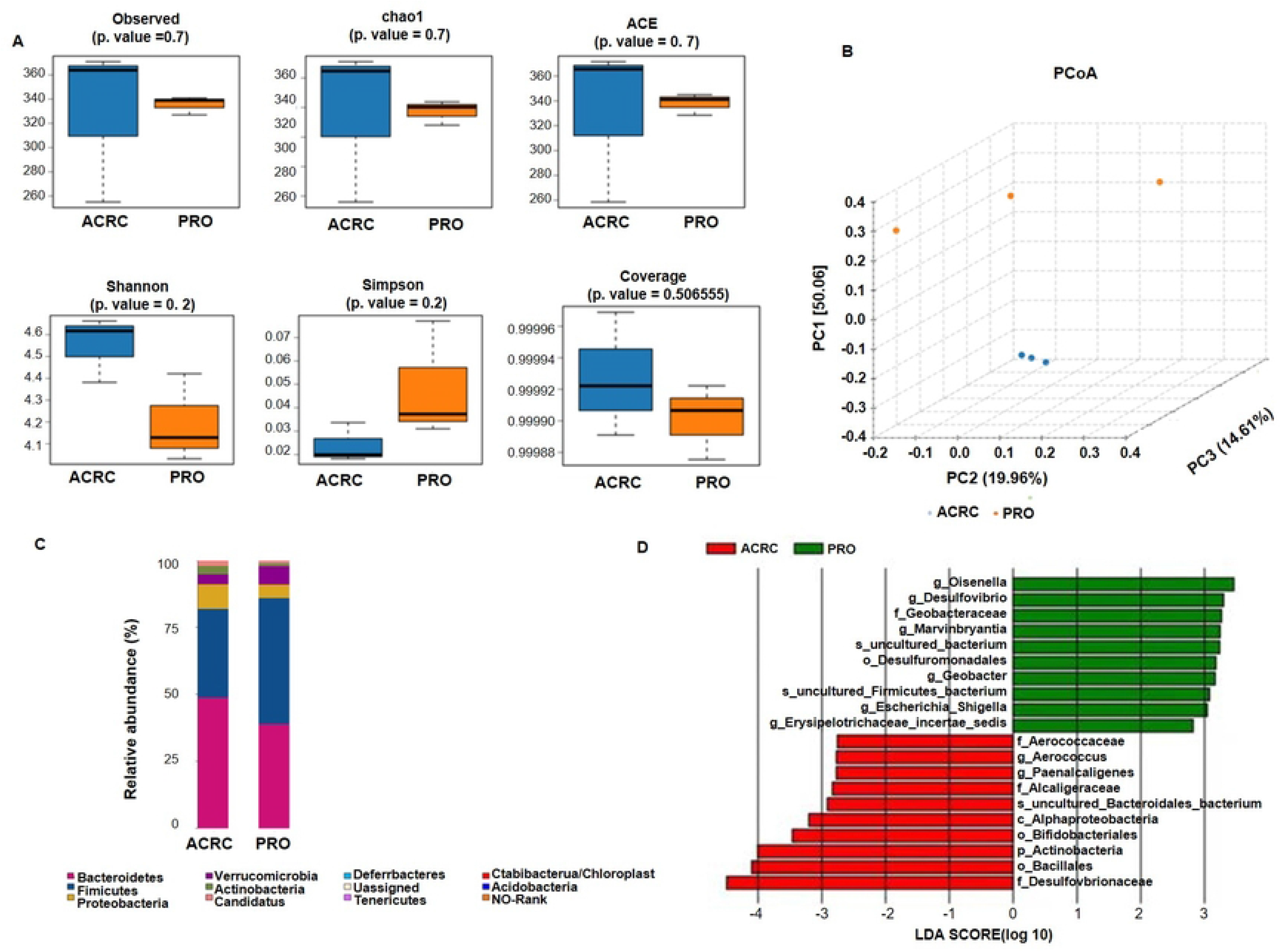
Intestinal microbial composition in PRO-treated CRC mice. (A) Alpha diversity analysis of stool samples. (B) PCoA analysis of stool samples. (C) Analysis of microbial species composition. (D) LEfSe analysis.

Significantly upregulated differential protein functional enrichment was found in nitroreductase, multidrug efflux pump subunit AcrA (membrane fusion protein), etc (Supplementary Figure 3B).

### FMT enhanced the therapeutic efficacy of OVV in advanced stage of CRC mice

We further evaluated the impact of FMT on the OVV-gp33 targeted therapy for advanced stage of CRC. Results showed the tumors grew at a slower rate in mice treated with FMT than in mice did not, and this tumor inhibition effect enhanced after treatment of OVV-gp33(Fig 7A, 7B). The tumor growth rate and tumor mass volume of mice received FMT was significantly slower than those of CRC mice that received probiotics, and this effect enhanced after treating with OVV-gp33 on the 17^th^ day (Fig 7C, 7D). The end event in CRC mice that received FMT and OVV-gp33 extended to 58 days (Fig 7E). This suggested that CRC may be more responsive to OVV-gp33 when combined with FMT than probiotics. FMT increased the abundance of activated CD8^+^ T cells and decreased the ratio of Treg cells in OVV-gp33-treated CRC mice, and FMT affected this antitumor immune response of OVV-gp33 to a much greater extent than probiotics (Fig 7F, 7G).

**Fig 7.**
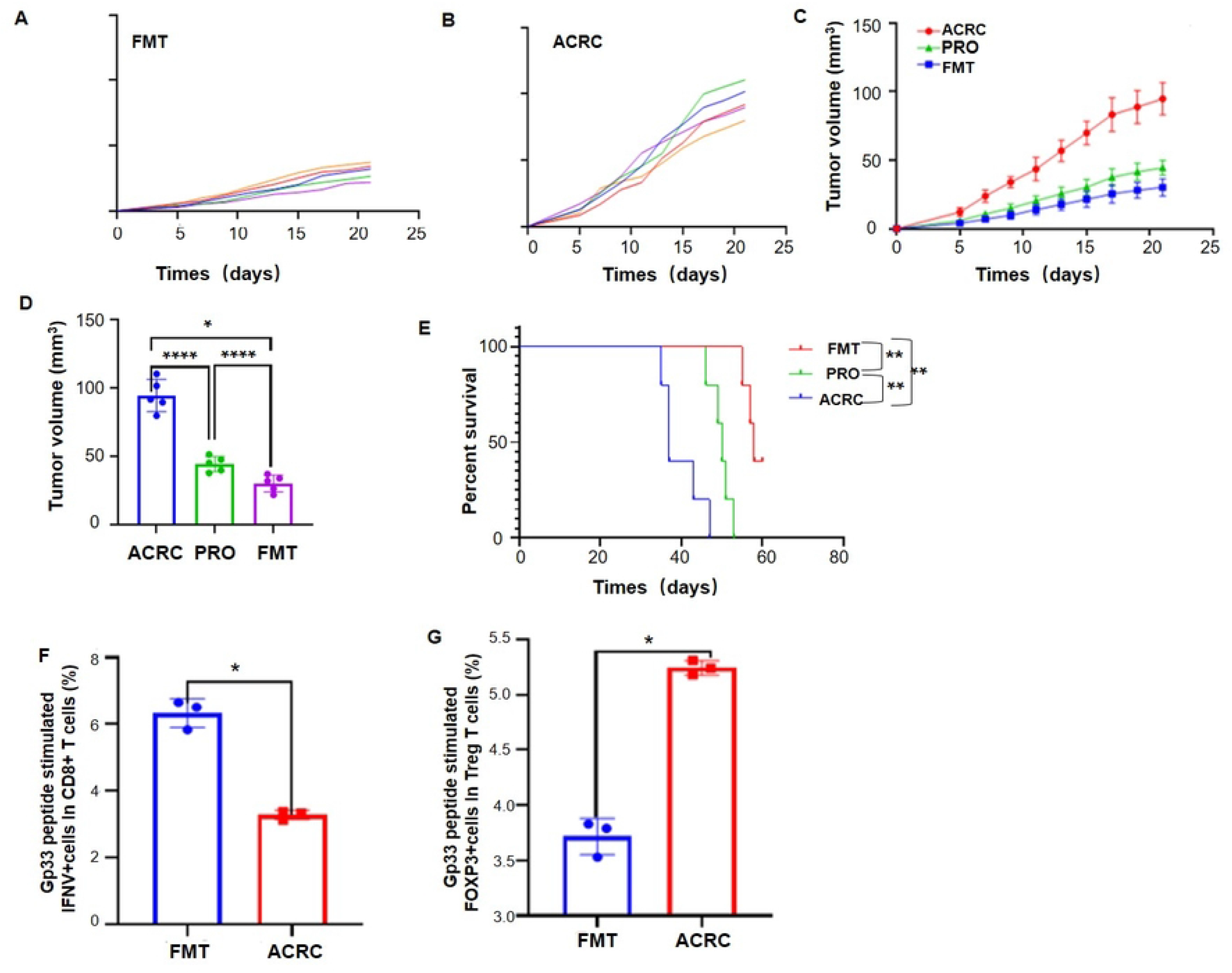
FMT improved the therapeutic effect of OVV-gp33 treatment. Tumor growth curve of FMT group (A) and ACRC group (B). (C), (D) Comparation of tumor volume among FMT group, PRO group and ACRC group. \*\*\*\**P*<0.05; \**P*< 0.01. (E) Survival curve of FMT group, PRO group and ACRC group. \*\**P*<0.05. (F) Activity of CD8^+^ T cells. \**P*<0.05. (G) Ratio of Treg cells. \**P*<0.05.

The abundance and diversity of the intestinal microbiota in CRC mice decreased when combined with FMT (Fig 8A), and beta diversity PCoA analysis also showed significant changes (Fig 8B). Bacteroidetes, Firmicutes, Proteobacteria and Verrucomicrobia were still dominant bacteria in mice that received FMT at the phylum level. The amount of Bacteroidetes and Proteobacteria decreased significantly, and Firmicutes and Verrucomicrobia increased significantly compared with mice in ACRC group, while Bacteroidetes and Actinomycetes increased compared with mice treated with probiotics (Fig 8C). The main enriched intestinal microbiota in mice that received FMT were Bacteroideaceae, *Oisenella*, *Coprobacter*, *Prevotella*, etc., which was different from mice that received probiotics (Fig 8D). The enrichment pathways of mice in FMT group were mainly vancomycin antibiotic biosynthesis, D alanine metabolism, secondary bile acid biosynthesis, glycolysis/gluconeogenesis, etc (Supplementary Fig 4A). The significantly down-regulated protein functions were mainly concentrated in DNA-binding transcription regulatory factors, LacI/PurR family, membrane proteins involved in the output of O antigen etc, and significantly upregulated differential protein functional enrichment was found in nitroreductase, outer membrane protein TolC, multidrug efflux pump subunit AcrA (membrane fusion protein), and nucleoside diphosphoglucoosinase (Supplementary Fig 4B).

**Fig 8.**
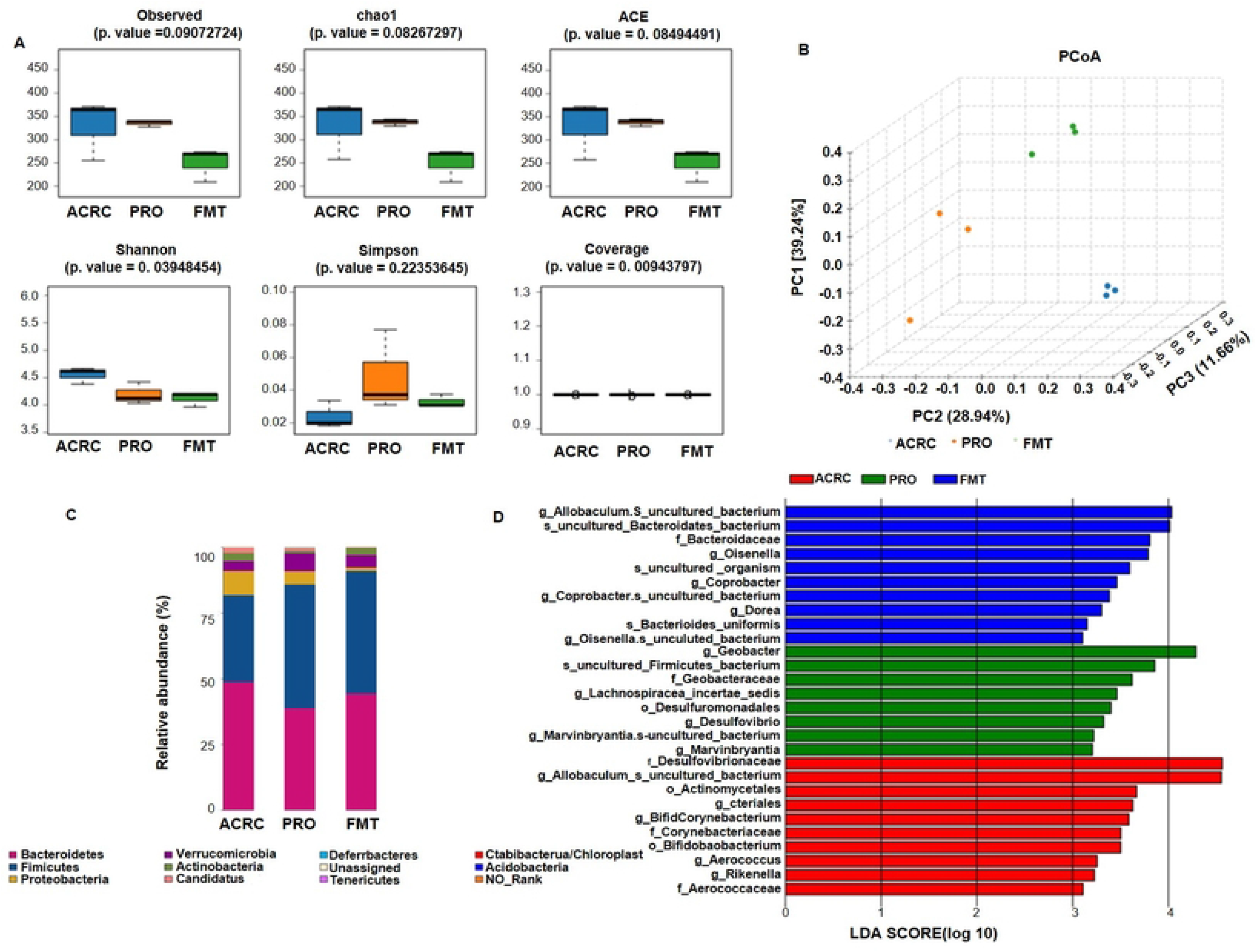
Intestinal microbial composition in FMT-treated CRC mice. (A) Alpha diversity analysis of stool samples. (B) PCoA analysis of stool samples. (C) Analysis of microbial species composition. (D) LEfSe analysis.

### Characteristics and variation trend of intestinal microbiota in CRC mice with different interventions

To gain further insight into the role of intestinal microbiota in OVV treatment in CRC, we summarized and analyzed the characteristics of microbiota in each group. Alpha diversity analysis showed that the total intestinal microbiota was increased after intradermal injection of MC38-gp33 cells, but decreased with the development of CRC. However, depletion with antibiotics significantly reduced the abundance and diversity of intestinal microbiota (Fig 9A). The microbial composition also varied in each group (Fig 9B). The analysis of microbial species composition showed that, at the phyla level, Bacteroidetes, Firmicutes, verrucobacteria and Proteobacteria were the dominant phyla in the microbial composition of mice except mice of ACRC group, while in ACRC group, Proteobacteria and verrucobacteria increased significantly, Bacteroidetes and Firmicutes decreased significantly (Fig 9C). There were 177 species of unique bacteria found in CON group, 268 species found in ECRC group, 179 species found in ACRC group, 55 species found in ABX group, 204 species found in PRO group, 110 species found in FMT group, and 7 species of common flora of all groups were found (Fig 9D). The intestinal flora in ECRC group were mainly enriched by *bifidobacteriaceae*, *actinomycetes*, *corynebacterium*, *rumen coccus* and *Aerococcus*. In ACRC group, Allobaculum, Bacillus, Barnesiella, Staphylococcus and enterorhabdus were dominant flora. The main enriched intestinal flora in ABX group were Proteobacteria, Enterobacterium, Parabacteroides, Morganella, *Citrobacter, Geobacteriaceae*, and *thiomonas*, etc. In PRO group, *Lactobacillus*, *Spirillaceae*, Spirillum and Firmicutes were dominant flora. The intestinal flora enriched in FMT group were *Erysipelotrichia*, *Ruatomella*, Prevotella, Euthrix and Bacteroides.

**Fig 9.**
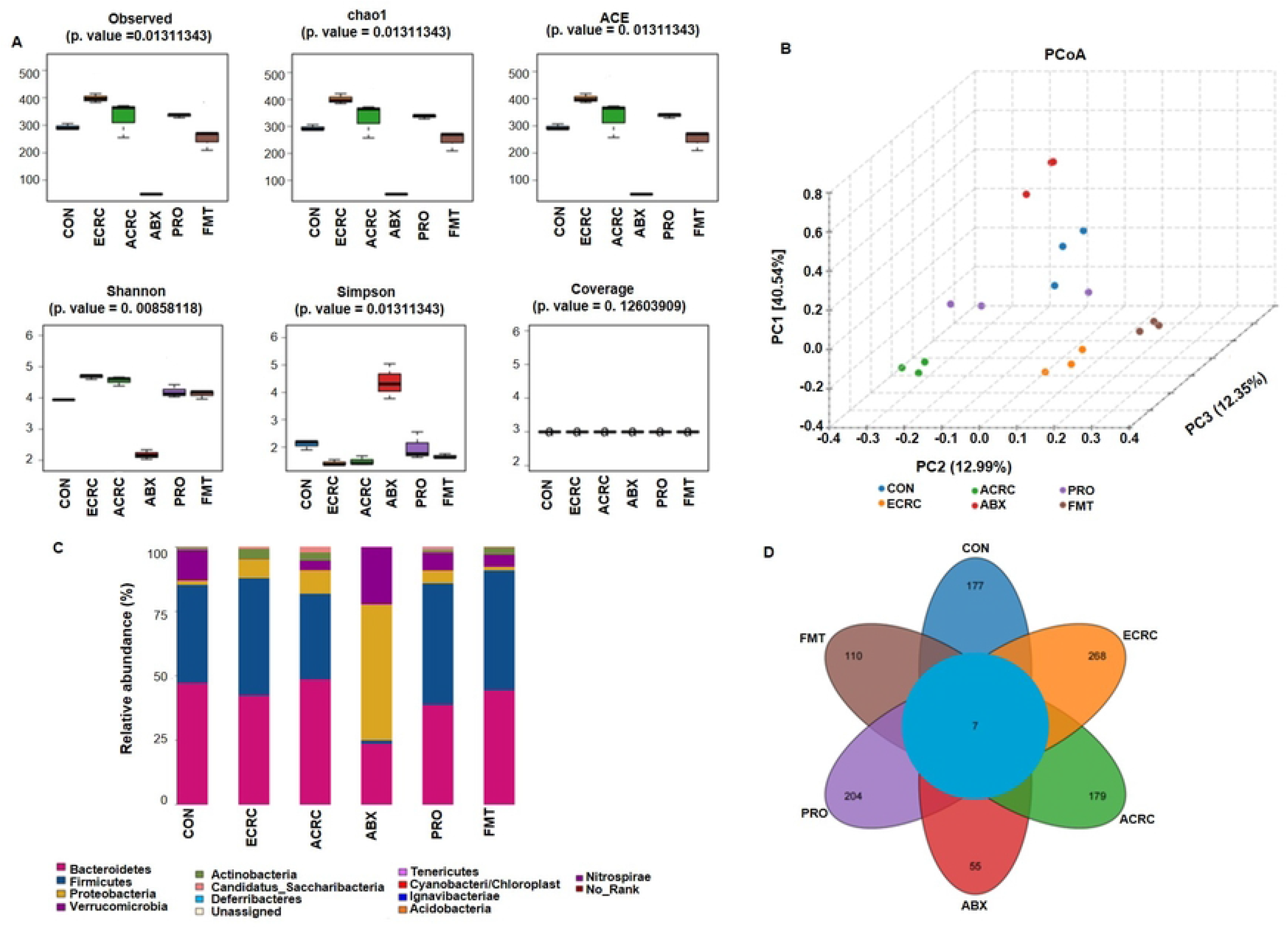
Characteristics and variation trend of intestinal microbiota in CRC mice of different groups. (A) Alpha diversity analysis of stool samples. (B) PCoA analysis of stool samples. (C) Analysis of microbial species composition. (D) Venn figure.

## Discussion

The present study provides new information on the combined treatment of intestinal microbiota and OVV in CRC. Specifically, we showed that OVV-gp33 treatment had a significant therapeutic effect on the early stage of CRC, but it was not as effective in the advanced stage of CRC. In addition, there were significant changes in the intestinal microbiota and T-cell activity in different stages of CRC. Furthermore, we found that altering the intestinal microbiota could modulate the therapeutic effect of OVV-gp33 and the subsequent T-cell response in advanced stages of CRC, confirming the role of the microbiota in tumor immunotherapy and providing a new combined therapy options for CRC.

OV is emerging as an alternative biological cancer therapy, which can selectively replicate and destroy tumor cells while leaving normal cells undamaged. OVV, a new Cancer vaccine, can target tumor-specific antigens and may even induce novel tumor-specific T cell responses[24]. The oncolytic effect to CRC was demonstrated in some OVs such as Newcastle disease virus[25], and recombinant Chinese measles virus vaccine strain Hu191[26]. These researches showed that OVs could induce apoptosis and inhibit cancer cell growth, but the inhibition effect in the different stages of CRC was not reported before. In the present study, the application of OVV-gp33 at Day 5 post tumor inoculation significantly inhibited tumor growth and improved the survival time of CRC mice, which is consistent with previous studies, but this effect was not obvious in mice treated with OVV-gp33 at Day 17 after tumor inoculation. We also found the activated CD8^+^ T cells decreased, Treg cells increased with the tumor progress. The level of CD4^+^ and CD8^+^ T cells in the systemic circulation is associated with the anti-tumor response of biologically targeted therapy, and this anti-tumor response maybe regulated by gut microbiome[27].

We therefore investigated the intestinal microbiota in different stages of CRC mice. With the development of CRC, the abundance and diversity of the intestinal microbiota decreased significantly, and its composition also changed. Bacteroidetes, Verrucobacteria and Proteobacteria increased, and Firmicutes and the ratio of Firmicutes to Bacteroidetes (F/B) decreased. Firmicutes and Bacteroidetes are two dominant phyla in gut microbiota, and the Firmicutes/Bacteroidetes ratio is widely believed to be related to gut homeostasis associated with some pathological conditions such as obesity and Type 2 Diabetes[28, 29]. Changes in intestinal microbial composition are an important causal factor in the development of CRC and can be indicative of cancer stage or disease progression [30, 31]. Therefore, the microbial profiles shift in present study maybe one of the causes which influence the therapeutic effect of OVV-gp33 treatment in advanced stage of CRC. Specific microbial signals induce the differentiation of T cells which is crucial for intestinal hemostasis[32]. Here, depletion of the intestinal microbiota resulted in impaired host immune responses, prevented CD8^+^ T-cell activation and increased ratio of Treg cells. Antibiotic therapy can reduce microbiome diversity and abundance, alter microbial species and then result in the dysregulation of host immune homeostasis[33]. Concurrently, disruption of the microbiota by ABX accelerated the tumor growth rate, shortened the survival time and eliminated the antitumor effect of OVV-gp33 treatment. Taken together, these data strongly suggested that the disruption of intestinal microbiota in the progression of CRC affects the activity of T cells, leading to the decline of immune response and the unsatisfied efficacy of OVV in CRC.

It is prospective that modulate intestinal microbiota can enhance efficacy of anti-tumor therapy[34]. We next treated MC38-gp33 bearing mice with probiotics and FMT respectively to change the microbiota composition in present study. The results showed that probiotics and FMT both enhanced the anti-tumor effect of OVV-gp33 treatment, and this tumor inhibition effect was more obvious in FMT- treated CRC mice. As revealed by 16S rDNA sequencing, the abundance and diversity of intestinal microbiota treated with probiotics or FMT were decreased, especially in FMT-treated mice with more Firmicutes and Verrucomicrobia. Intestinal microbiota of FMT treated mice were mainly enriched in Bacteroideae, Eurosenella, Coprobacter, Prevotella, etc. Prevotella is reported to be associated with augmented T helper type 17 (Th17) -mediated mucosal inflammation and can promote inflammatory disorders[35]. Previous study reported that intestinal microbiota stimulates chemokine production by CRC cells, thus promoting the recruitment of beneficial T cells into tumor tissues[36]. Similarly, OV exhibits its antitumor role mainly by amplifying particular anticancer immune reactions through T-cell activation[37]. Intestinal microbiota can influence cancer immunotherapy through the host’s immune system[38], promote the ability of immunotherapy to raise endogenous tumor-reactive T cell responses[39]. In present study, probiotics and FMT changed the composition and diversity of intestinal microbiota and modulate the T cells subsets, which then restored the immunotherapy response of OVV-gp33. Therefore, these results suggested that microbiota can be applied as a complement of OVV therapy to achieve better anti-tumor outcomes in CRC.

Additionally, we found that there were several common metabolic pathways in ECRC group, PRO group and FMT group, mainly glycolysis/gluconeogenesis, pentose phosphate pathway and lysine biosynthesis pathway. However, the enrichment pathways in mice of ACRC group and ABX group were more inclined to the carbohydrate, secondary bile acid biosynthesis, amino acid, biotin metabolism. It is proposed that microbial metabolic alteration is correlated with CRC progression [40], such as secondary bile acids, which is a potentially carcinogenic metabolite[41].On the other hand, FMT, probiotics that restore the intestinal microbial can promote the activation of T cells by enhancing the glycolysis/gluconeogenesis, pentose phosphate pathway and lysine biosynthesis pathway. Increased glycolysis is considered a hallmark metabolic change in most immune cells undergoing rapid activation, is crucial for the functioning of T helper and peripherally Treg cells[42]. The pentose phosphate pathway is a branch of glycolysis that produces ribose for nucleotides used in DNA and RNA synthesis, or glutathione biosynthesis that promotes antioxidant reactions[43]. Lysine is considered to be a nonspecific bridging molecule that promotes immunoglobulin production and lymphocyte proliferation to supply the anabolic process of the adaptive response[44]. Therefore, we believe that the link between these metabolic pathways and the activation of T cells contributes to the immunotherapeutic efficacy of OVV-gp33 in CRC.

## Conclusions

In summary, the obtained data indicate that OVV-gp33 is well suited to the early stage of CRC but is not as effective in the advanced stage due to the intestinal microbial composition difference. However, the combination of probiotics or FMT could enhance the antitumor effect of OVV-gp33 in advanced stage of CRC by modulating the microbial communities, metabolites and altering T-cell activity. Our study may provide a promising combination strategy of microbial modification and OVV-targeted antigen therapy in CRC.

## Author contributions

Writing-original draft: Xia Chen.

Formal analysis: Lin Qin.

Data curation: Guang-Jun Wang.

Conceptualization: Bing Hu.

Conceptualization, Supervision: Jun Li.

## Acknowledgments

None

## Data Availability

All relevant data are within the manuscript and its Supporting Information files.

